# Environmental factors dominate over host genetics in shaping human gut microbiota composition

**DOI:** 10.1101/150540

**Authors:** Daphna Rothschild, Omer Weissbrod, Elad Barkan, Tal Korem, David Zeevi, Paul I Costea, Anastasia Godneva, Iris Kalka, Noam Bar, Niv Zmora, Meirav Pevsner-Fischer, David Israeli, Noa Kosower, Gal Malka, Bat Chen Wolf, Tali Avnit-Sagi, Maya Lotan-Pompan, Adina Weinberger, Zamir Halpern, Shai Carmi, Eran Elinav, Eran Segal

## Abstract

Human gut microbiome composition is shaped by multiple host intrinsic and extrinsic factors, but the relative contribution of host genetic compared to environmental factors remains elusive. Here, we genotyped a cohort of 696 healthy individuals from several distinct ancestral origins and a relatively common environment, and demonstrate that there is no statistically significant association between microbiome composition and ethnicity, single nucleotide polymorphisms (SNPs), or overall genetic similarity, and that only 5 of 211 (2.4%) previously reported microbiome-SNP associations replicate in our cohort. In contrast, we find similarities in the microbiome composition of genetically unrelated individuals who share a household. We define the term *biome-explainability* as the variance of a host phenotype explained by the microbiome after accounting for the contribution of human genetics. Consistent with our finding that microbiome and host genetics are largely independent, we find significant biome-explainability levels of 16-33% for body mass index (BMI), fasting glucose, high-density lipoprotein (HDL) cholesterol, waist circumference, waist-hip ratio (WHR), and lactose consumption. We further show that several human phenotypes can be predicted substantially more accurately when adding microbiome data to host genetics data, and that the contribution of both data sources to prediction accuracy is largely additive. Overall, our results suggest that human microbiome composition is dominated by environmental factors rather than by host genetics.

## Introduction

The gut microbiome is increasingly recognized as having fundamental roles in multiple aspects of human physiology and health including obesity, non-alcoholic fatty liver disease, inflammatory diseases, cancer, metabolic diseases, aging, and neurodegenerative disorders^1–14^. Humans acquire bacteria at birth and are exposed to bacteria from the environment throughout their lifespan^15–18^. Although mammalian microbiome composition may change during life^19^, it is considered relatively stable^20,21^.

A fundamental question is the extent to which microbiome composition is determined by host genetics as opposed to being shaped by the environment. Previous studies identified several heritable bacterial taxa and SNPs associated with the gut microbiome, but the effect of host genetics on the overall microbiome composition has not been investigated. For example, a study of twins identified 33 significantly heritable bacterial taxa^22,23^, but did not evaluate the combined relative abundance of these taxa. Another study identified several bacteria taxa shared among non-twin family members^24^, but could not distinguish between genetic and environmental factors, because teasing these two factors apart is typically only done with twin studies^25^.

Several recent studies tested for association between host SNPs and individual taxa or pathways^24,26–28^. However, a major statistical challenge in such studies is the large number of hypotheses tested, since there are millions of human SNPs and thousands of bacterial taxa and pathways^29–33^. Consequently, most of the associations reported in these studies are not statistically significant after multiple hypothesis testing correction. To alleviate the multiple hypothesis burden that results from examining associations to individual bacteria, one study tested for associations between each SNP and gut microbiome β-diversity, and found 42 significant loci that together explain 10% of the variance of the β-diversity^27^. However, this study did not provide an assessment of the statistical significance of this result. Thus, given the overall limited number of significant findings reported to date, the extent to which human genetics shapes microbiome composition remains unclear.

In this work, we studied microbial-genetic associations using a cohort of 696 healthy Israeli individuals for whom we obtained information on genotypes, metagenome and 16S-sequenced gut microbiomes, numerous anthropometric and blood phenotypes, and dietary habits^34^. Notably, individuals in our cohort represent several different ancestral origins yet they share a relatively homogeneous environment, since most Israeli individuals lead similar ‘Western’ life styles.

Our results demonstrate that environmental factors have a substantially stronger effect on microbiome composition than host genetics, suggesting that microbiome composition is predominantly shaped by environmental factors. Specifically, we show that there is no statistically significant association between microbiome composition and individual SNPs, genetic ancestry, or overall genetic similarity. Notably, our study is well powered to find microbiome associations with ancestry or genetic similarity, as such tests do not suffer from a multiple hypothesis burden. We further show that family relatives with no history of a shared household do not have similar microbiomes, whereas we find microbiome similarity among genetically unrelated individuals who share a household.

As our findings suggest that microbiome and host genetics are largely independent, we aimed to disentangle and compare the association of human phenotypes with microbiome and with host genetics. Several bacterial taxa have been shown to explain a significant fraction of the variance of BMI, HDL cholesterol, and triglycerides, after accounting for host genetics^35^, but the association of the overall microbiome composition with these and other phenotypes has not been quantified to date. We define the term *biome-explainability*, which analogously to genetic heritability quantifies the overall association between the microbiome and host phenotypes after accounting for the association of host genetics. We find significant biome-explainability levels of 16-33% for BMI, fasting glucose, glycemic status, HDL cholesterol, waist circumference, waist-hip ratio, and lactose consumption. Finally, we demonstrate that adding microbiome data on top of human genetics substantially improves human phenotype prediction accuracy and that the contribution of both data sources to the accuracy is mostly additive. These results suggest that the microbiome should be routinely considered in addition to genetics in studies aimed at explaining the variance of human phenotypes.

## Results

### Cohort and analysis description

We studied a cohort of 696 healthy Israeli adults for whom we collected for every participant blood for genotyping and phenotyping, stool for metagenome and 16S rRNA-gene sequencing, anthropometric measurements, and answers to a food frequency questionnaire^34^ (**Table 1**). We performed genotyping at 712,540 SNPs and imputed them to 5,578,121 SNPs (Methods). Stool samples were profiled using both metagenome and 16S rRNA-gene sequencing. Metagenomes were subsampled to 10M reads per sample to achieve even sequencing depth across individuals.

**Table 1.**
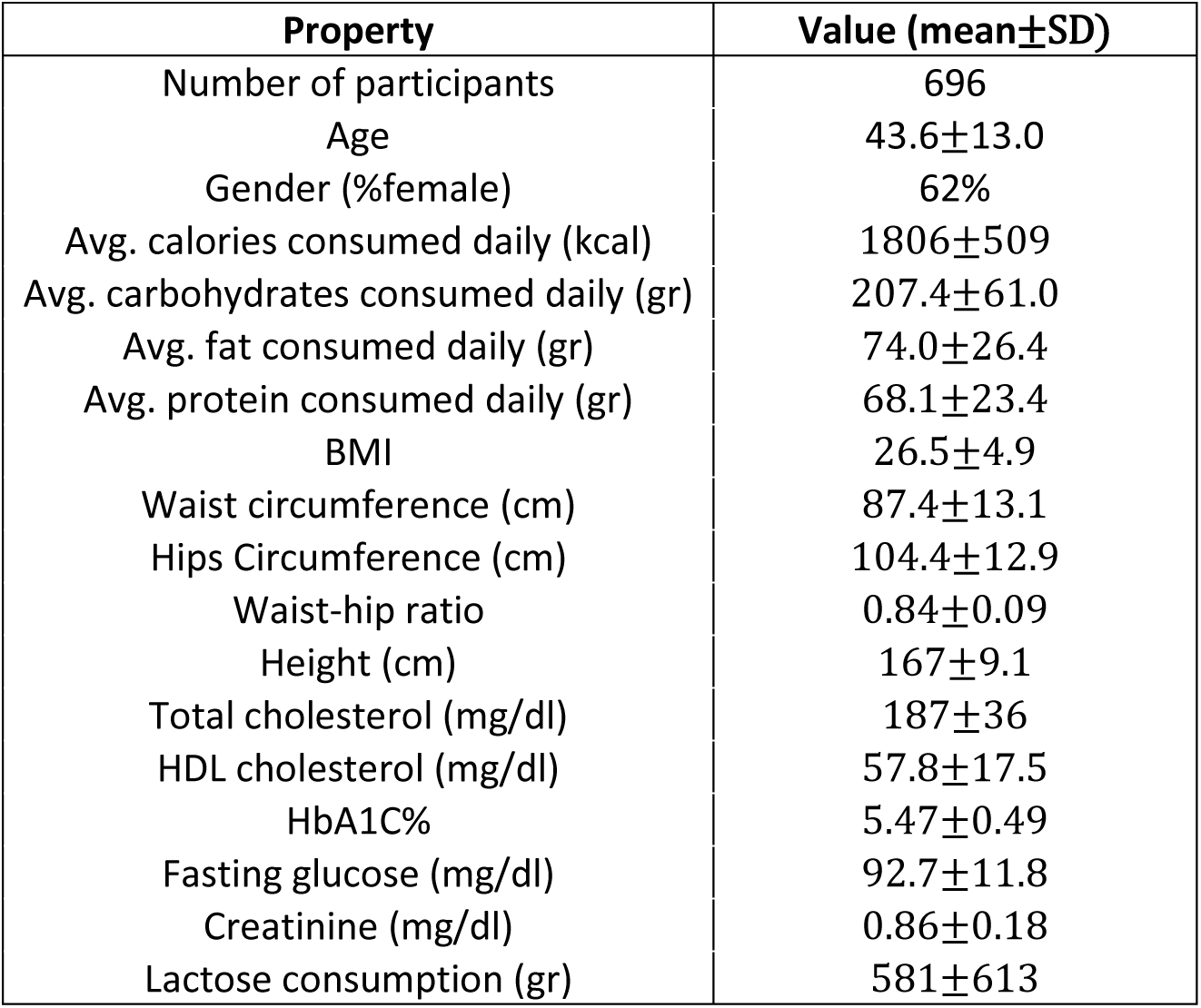
Baseline characteristics of the cohort. Shown are the mean and standard deviation of all properties used either as covariates or as investigated phenotypes. Dietary properties are based on information recorded in real time by study participants on their smartphones (see Methods).

| Property | Value (mean±SD) |
| --- | --- |
| Number of participants | 696 |
| Age | 43.6±13.0 |
| Gender (%female) | 62% |
| Avg. calories consumed daily (kcal) | 1806±509 |
| Avg. carbohydrates consumed daily (gr) | 207.4±61.0 |
| Avg. fat consumed daily (gr) | 74.0±26.4 |
| Avg. protein consumed daily (gr) | 68.1±23.4 |
| BMI | 26.5±4.9 |
| Waist circumference (cm) | 87.4±13.1 |
| Hips Circumference (cm) | 104.4±12.9 |
| Waist-hip ratio | 0.84±0.09 |
| Height (cm) | 167±9.1 |
| Total cholesterol (mg/dl) | 187±36 |
| HDL cholesterol (mg/dl) | 57.8±17.5 |
| HbA1C% | 5.47±0.49 |
| Fasting glucose (mg/dl) | 92.7±11.8 |
| Creatinine (mg/dl) | 0.86±0.18 |
| Lactose consumption (gr) | 581±613 |

Unless stated otherwise, we excluded a subset of the individuals so that no pair of individuals are close relatives (having a fifth or stronger degree of relation) or share a household (Methods). Finally, we included covariates for age, gender, stool collection method, and self-reported daily average caloric, fat, protein and carbohydrates consumption (Methods). When not examining association of bacteria with ancestry or kinship, we also included the top five principal components (PCs) of the genotypes as covariates.

### Microbiome composition is not associated with ancestry or genetic kinship

Our population is one of the largest cohorts of diverse Jewish individuals genotyped in a single study, consisting of self-reported Ashkenazi (n=346), North African (38), Middle Eastern (23), Sephardi (8), Yemenite (7), admixed (243) and unknown/other (31) ancestries (Methods). Genetic ancestry is often reflected in the top principal components (PCs) of the genotypes^36^. We therefore computed the PCs via PC-AiR^37^, which is robust to relatedness and admixture (Methods), and verified that there is a close correspondence between the top two PCs and self-reported ancestry of individuals with a single ancestral origin (P<10^-21^ for both PC1 and PC2, Kruskal-Wallis test; **Fig. 1a**, **Table 2 column 1 row 1**, **Supplementary Table 1**).

**Figure 1:**
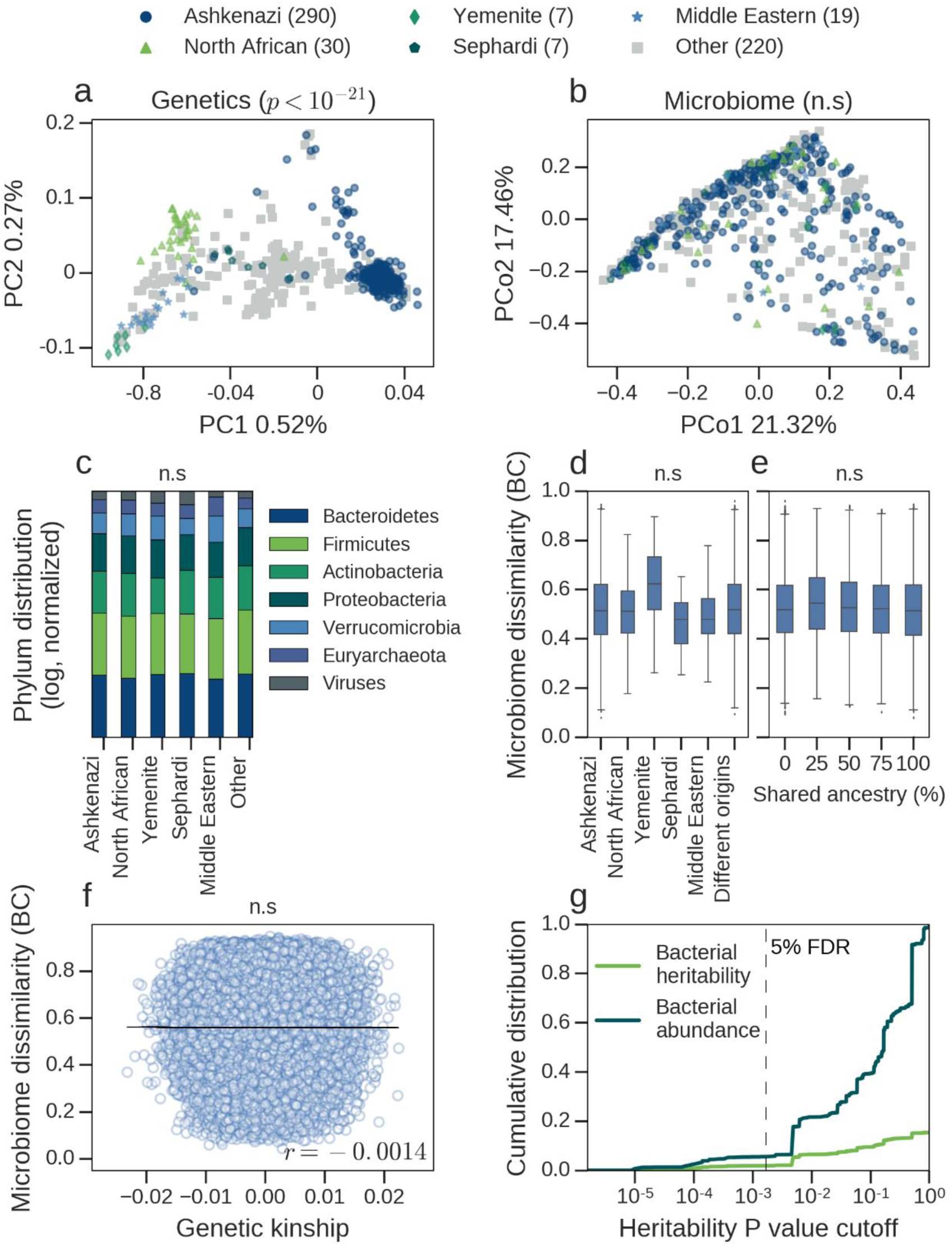
Genetic similarity is not associated with similarity in microbiome composition. **(a)** Principal component analysis of host genotypes. Markers represent individuals, with self-reported ancestry being represented by different colored shapes. Admixed individuals and individuals with a partly unknown origin are assigned to the group *Other*. Note the clear separation of individuals by genetic ancestry, with the top two PCs significantly associated with genetic ancestry (Kruskal-Wallis test). **(b)** Same as (a), but for principal coordinate analysis of the microbiome at the genus level, based on Bray-Curtis dissimilarity. Note that here individuals do not separate by ancestry, and no significant association is detected (Kruskal-Wallis test). **(c)** Distribution of average phylum abundance among individuals with a single ancestral origin (in log scale, normalized to sum to 1.0), for each phylum with an average study-wide relative abundance >0.1%. There is no significant difference in phyla distribution across different ancestral origins (Kruskal-Wallis test for each phylum). **(d)** Box plots showing the distribution of Bray-Curtis dissimilarities at the genus level across pairs of individuals with a single ancestral origin (first five box plots), and across pairs with a single but different ancestral origin (right box plot). The numbers of pairs are Ashkenazi (n=41,905), North African (n=435), Yemenite (n=21), Sephardi (n=21), Middle Eastern (n=171) and pairs with different origins (n=57,575). The markers represent the (5%, 95%) percentiles of the distribution. There is no significant difference in Bray-Curtis dissimilarities across the different groups (Kruskal-Wallis test for the top five Bray-Curtis PCos). **(e)** Box plots showing the distribution of Bray-Curtis dissimilarities at the genus level across pairs of individuals, organized according to shared ancestry fraction (the fraction of grandparents born in countries associated with the same ancestry), for pairs with 0% (n=11,4254), 25% (n=23,880), 50% (n=77,142), 75% (n=24,430) and 100% (n=88,623) shared ancestry fractions. Pairs of individuals who are more ancestrally similar do not have significantly more similar microbiomes (Mantel test). **(f)** Genetic kinship values (x axis) versus Bray-Curtis dissimilarities at the genus level (y axis) between pairs of individuals. Markers represent pairs of individuals. The black line is the regression slope, and the Pearson correlation (r) is displayed. Pairs of individuals who are more genetically similar do not have significantly more similar microbiomes (Mantel test). **(g)** Analysis of the overall heritability of the microbiome, based on bacterial heritability estimates reported in the twins study of Goodrich *et al.*^23^. The x axis represents P values of heritability estimates of bacterial taxa, as reported in Goodrich *et al.* The y axis represents (1) cumulative bacterial abundance, whose value at any point *k* along the x-axis is the sum of relative abundances of all taxa with P<*k* (dark curve); and (2) cumulative estimated microbiome heritability, whose value at any point *k* along the x-axis is the sum of heritability estimates of all taxa with P<*k*, weighted by their relative abundance (light curve). When P corresponds to a 5% FDR, the overall microbiome heritability is 1.9%.

**Table 2:** No significant association between ancestral or genetic similarity and the gut microbiome. Each cell contains the P value of a single or multiple statistical tests, testing if individuals who are more similar according to ancestry or genetic kinship (in columns) are also more similar according to (1) microbiome β-diversity (using Bray-Curtis dissimilarity); (2) microbiome a-diversity (using Shannon diversity); (3) abundance of specific taxa; or (4) genetic kinship (in rows). The first column includes n=353 individuals with a single ancestral origin. The second and third columns include n=573 individuals. P values in the first column are based on Kruskal-Wallis tests (using the top 5 microbiome PCos for Bray-Curtis dissimilarity, and the top 5 genetic PCs for genetic kinship); P values in the other columns are based on Mantel tests (using Euclidean distances for ancestry proportions; see Methods).

|  |  | Ancestry<br>(categorical) | Ancestry<br>(proportions) | Genetic kinship |
| --- | --- | --- | --- | --- |
| Host genetics<br>(control) | Kinship | <0.03 | <10<sup>-5</sup> | - |
| Microbiome | β-diversity | >0.13 | 0.84 | 0.58 |
|  | α-diversity | 0.25 | 0.92 | 0.67 |
|  | Specific taxa | >0.05 | >0.05 | >0.05 |

In contrast to the association between ancestry and genetics, we found no significant association between ancestry and microbiome composition. Specifically, there was no significant correlation between any of the top five host genetic PCs and any of the top five microbiome β-diversity principal coordinates (computed using Bray-Curtis dissimilarity at the genus level; P>0.16 for all pairwise associations, Spearman correlation; **Supplementary Table 2**). We next tested if individuals sharing an ancestry have a more similar (1) microbiome composition (quantified by PCos of Bray-Curtis dissimilarities); (2) microbiome α-diversity (quantified by Shannon diversity index^38^); and (3) abundance of specific taxa. In all cases, we found no significant association (Kruskal-Wallis test; **Fig. 1b**-**c**, **Table 2 column 1**, **Supplementary Table 1**).

The lack of microbiome-ancestry associations does not rule out the possibility that individuals who are more ancestrally or genetically similar have more similar microbiomes. To test this hypothesis, we first constructed an ancestral similarity matrix (using Euclidean distances of ancestry proportions; Methods) and a genetic similarity matrix (using genetic kinship; Methods) for all pairs of individuals, including individuals with more than one ancestral origin. We then tested if pairs of individuals who are more ancestrally or genetically similar have a more similar (1) microbiome composition (quantified by Bray-Curtis dissimilarity at the genus level); (2) a-diversity; and (3) abundance of specific taxa. As before, in all cases we found no significant association (Mantel test^39^; Methods, **Fig. 1d-f, Table 2 columns 2-3**, **Supplementary Tables 3-4**). As a control, we verified that ancestrally similar individuals are significantly similar genetically (P<10^-5^, Mantel test; **Table 2 column 2 row 1**, **Supplementary Table 3**).

We also applied several different machine learning prediction algorithms to try and predict ancestry proportions from microbiome but none were successful (prediction R^2^<0.01 for all ancestries; Methods). Additionally, canonical correlation analysis^40^, a well-known technique for testing for low rank linear dependencies between high dimensional objects, showed no association between host genotypes and microbiome abundances (P=0.21, permutation testing; Methods, **Supplementary Table 5**).

While the above results are based on metagenome-derived genus level abundances after regressing out covariates, we obtained similar results also when using any of the following: Other metagenome-derived taxonomic and functional levels (phylum, class, order, family, species, and bacterial genes; see Methods); 16S data; non-metric multidimensional scaling^41^ instead of PCoA; and when omitting covariates (**Supplementary Figs. 1-3, Supplementary Tables 1-5**).

Finally, we asked whether our results are in line with data from previous studies showing that several bacterial taxa are significantly heritable^23,24,26–28^, by analyzing data from the twins study of Goodrich *et al*^23^. First, we found that the sum of the relative abundances of all 33 taxa reported as significantly heritable in Goodrich *et al.*^23^ is only 5.6% (Methods). Next, we found that the combined heritability of significantly heritable taxa in this data set (weighted by their relative abundance) was only 1.9%, or at most 8.1% when not correcting for multiple comparisons (Methods, **Figure 1g, Supplementary Table 6**). These numbers can serve as estimates of the lower and upper bound of the true overall microbiome heritability. These results are thus consistent with the results obtained on our cohort and collectively, they suggest that host genetics is not a major determinant of gut microbiome composition.

### Microbiome β-diversity is not associated with individual SNPs

We next tested for associations between individual SNPs and microbiome β-diversity, by testing if individuals with a smaller Bray-Curtis dissimilarity are more likely to share the same allele via MiRKAT^42^ (Methods), and found no significantly associated SNPs (**Fig. 2a**).

**Figure 2:**
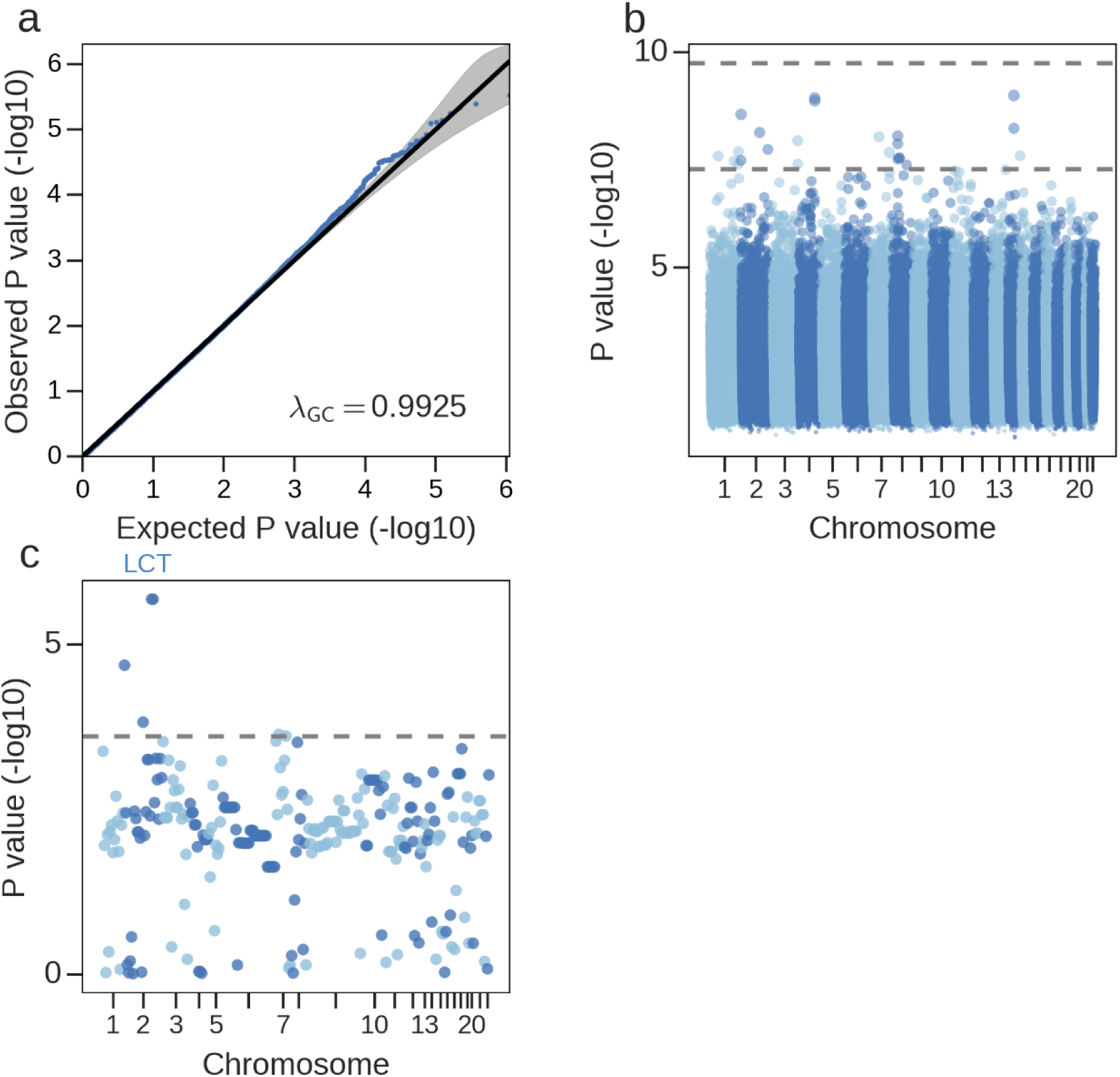
Limited evidence for microbiome associations with specific SNPs. **(a)** A quantile-quantile (qq) plot of a genome-wide test which tests every genotyped SNP for association with microbiome β-diversity, by examining whether individuals with a smaller Bray-Curtis dissimilarity tend to carry more similar alleles at this SNP (using a linear mixed model based on Bray Curtis dissimilarities at the genus level, with n=573 individuals; see Methods). Markers represent obtained P values (y axis) versus their expected value under the null hypothesis of no association (x axis). No SNP is significantly associated with the microbiome β-diversity at a level of 5% FDR. λ_GC_ is the genomic control inflation factor^95^, where deviation from 1.0 indicates an inflation or deflation of P values. **(b)** A Manhattan plot showing the lowest p-value obtained for every SNP tested for association with 288 taxa and with microbiome β-diversity, using n=665 individuals. The dashed lines represent a genome-wide significant P value for a single tested taxon (5×10^−8^), and corrected for testing 288 different taxa (5×10^−8^ / 288). **(c)** A Manhattan plot with the lowest P-value obtained for 225 SNPs in 211 loci previously reported in one or several previous studies to be significantly microbiome-associated^23,24,26–28^, using n=665 individuals. The dashed line represents the P value for successful replication (0.05 / 211). Five SNPs are successfully replicated (rs4988235, rs6730157, rs7581129, rs1360741, rs7801810), with two SNPs (rs4988235 and rs6730157) residing in close vicinity to the LCT gene.

In addition, our data did not replicate significant associations for any of the 42 SNPs reported by Wang et al. to be significantly associated with microbiome β-diversity^27^ (P>0.05 for all previously reported SNPs). To verify that this is not due to the different testing methods used, we also tested all SNPs in our cohort using the method of Wang *et al.* and again could not replicate these SNPs (P>0.05 for all SNPs; Methods).

Wang *et al.* also reported that their 42 reported SNPs accounted for 10% of the β-diversity, but did not report statistical significance for this result. To assess the potential significance of such a result, we first applied the method of Wang *et al.* to our data and found that the 42 top ranked SNPs accounted for 13.5% of the β-diversity. We then estimated the significance of this result by randomly permuting the microbiome assignment of individuals in our data 100 times, so that every individual has the microbiome of a randomly selected individual. In each permutation, we ranked all SNPs, and explained the β-diversity variance using the 42 top ranked SNPs (which were all top ranked by chance due to the random permutations; Methods). The explained variance fractions across the permutations ranged from 11.2% to 20.0%, with 85% of the results being greater than the one obtained with non-permuted data, thus yielding P=0.85. We conclude that accounting for >10% of the microbiome composition via top ranked SNPs may be an inherent property of the method employed and not a biologically meaningful result.

Thus, we find no evidence in our data for association of any individual SNP with microbiome β-diversity, including for SNPs previously reported to exhibit such association.

### Limited evidence for SNP associations with specific taxa

We next tested for association between individual SNPs and specific taxa. Due to the large number of tested hypotheses, we maximized power by including pairs of individuals with a shared household or with close genetic relatedness, while appropriately controlling for these potential cofounding sources via a linear mixed model (LMM) (Methods). We dichotomized taxa with a zero-inflated distribution (**Supplementary Table 7**) and only tested for presence/absence patterns for these bacteria, in order to alleviate modeling violations (Methods). This analysis identified 167 SNPs with P<5×10^−8^, corresponding to a false discovery rate (FDR) of 29%, but none remained statistically significant at an FDR of 5% (**Fig. 2b**, **Supplementary Table 8**).We next examined the association of 225 SNPs in 211 genetic loci reported as significantly associated with specific taxa or with β-diversity in any of five previous studies^23,24,26–28^ (Methods). To maximize replication power, we used the minimal P value obtained for each SNP across all taxa belonging to the same phylum. Only 5 of the 211 loci (2.5%) replicated at P < 0.05/211 (**Fig. 2c, Supplementary Table 9**; Methods). Two of these 5 loci reside in close vicinity to the LCT gene, and were found by several previous studies to be associated with the Bifidobacterium genus, possibly owing to its association with lactose consumption^43,44^.

Notably, the LCT gene is the only case in which there was an overlap between the SNPs reported in any pair of five previous studies. Moreover, no pair of previously reported SNPs from any two studies were within 100Kb of each other, or within 1Mb of each other and associated with taxa belonging to the same phylum (**Supplementary Table 9**).

### Microbiome composition is not associated with familial relations but is moderately associated with household sharing

We next asked whether family members with no history of household sharing have similar microbiome compositions. To this end, we extracted 11 pairs of individuals from our data whose kinship coefficient was between the standard cutoffs for 2^nd^ degree and 5^th^ degree relatives^45^ and who do not share a household. We then asked whether their average Bray-Curtis dissimilarity is significantly smaller than across non-related pairs who do not share a household via permutation testing (Methods), and found no such evidence at any taxonomic level or at the level of bacterial gene abundance (P>0.9 for phylum, class, order, family, genus, species, and bacterial gene abundance; **Fig. 3**, **Supplementary Table 10**). In contrast, we found borderline significant microbiome sharing in 22 first-degree relative pairs, who are likely to have a history of household sharing, at the phylum (P=0.033) and order (P=0.006) taxonomic levels, but no significant sharing at other levels (**Fig. 3, Supplementary Table 10**).

**Figure 3:**
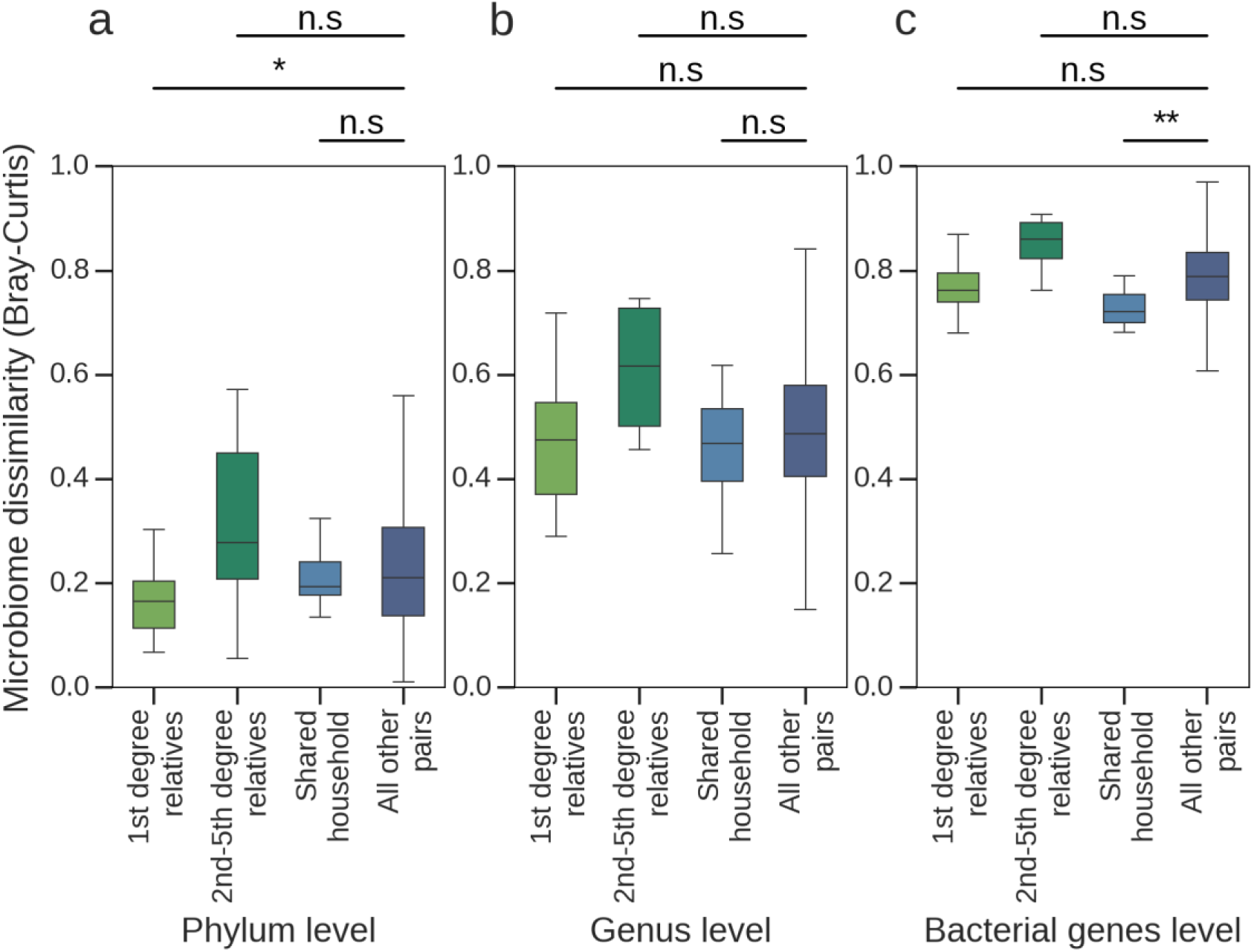
Individuals who share a household at present or in the past have significantly correlated microbiomes. Shown are box plots depicting the distribution of Bray-Curtis dissimilarities across pairs of individuals at the **(a)** phylum; **(b)** genus; and **(c)** bacterial genes level. Each panel shows the Bray-Curtis dissimilaries among all pairs of (1) individuals who are 1st degree relatives, and hence likely to once have shared a household (n=22 pairs); (2) individuals who are 2nd, 3rd, 4th or 5th degree relatives, and hence unlikely to have a present or past shared household (n=11 pairs); (3) individuals self-reported to currently share a household (n=12 pairs); and (4) all other pairs of individuals (n=53,923 pairs). P values were computed via 100,000 permutation tests. First degree relatives have significantly similar bacterial phyla and bacterial gene abundances, and individuals with present household sharing have significantly similar bacterial gene abundances. * P<0.05; ** P<0.005.

To test the effect of present household sharing, we repeated the above analysis for 12 pairs of genetically unrelated individuals who reported living in the same household during data collection. Although we did not detect microbiome sharing at any taxonomic level, we did find statistically significant microbiome sharing at the level of bacterial gene abundance (P=0.004; **Fig. 3, Supplementary Table 10**).

Thus, these results suggest that past or present household sharing may partly determine gut microbiome composition, whereas we find no supporting evidence for microbiome sharing among family relatives with no past household sharing. While these results do not rule out the possibility of specific heritable bacterial taxa, it suggests that such taxa likely compose a small portion of the microbiome, with a minor effect on overall microbiome sharing. Our results corroborate those of Goodrich *et al.*^22^, who showed using larger samples than ours that twins have significantly correlated microbiomes compared to non-related individuals (P<0.009), and that microbiome similarity among monozygotic twins compared to dizygotic twins is only borderline significant (P=0.032 under an unweighted UniFrac dissimilarity, P>0.05 under Bray-Curtis and weighted UniFrac dissimilarity).

### Biome-explainability can be assessed more accurately than genetic heritability

Given the lack of evidence for strong genetic-microbiome associations, we next asked how well different host phenotypes can be inferred based on the microbiome, as compared to host genetics. In statistical genetics, the fraction of phenotypic variance explained by genetic factors is called heritability, and is typically evaluated under an LMM framework^46^. Intuitively, LMMs estimate heritability by measuring the correlation between the genetic kinship and the phenotypic similarity of pairs of individuals. We define the analogous term of *biome-explainability*, which corresponds to the fraction of phenotypic variance explained by the microbiome. Specifically, we construct the microbiome “kinship” matrix as a microbiome-similarity matrix based on presence-absence patterns of bacterial genes extracted from metagenomic samples (Methods).

Genetic heritability estimation requires samples with thousands of individuals to yield reliable estimates^47^. We therefore first asked whether biome-explainability can be reliably estimated with our sample size. To this end, we computed confidence intervals (CIs) for a large number of biome-explainability values via parametric bootstrap, which constructs CIs by repeatedly drawing random realizations of the phenotypes according to the principle of test inversion^48^. We find that biome-explainability can be estimated more accurately than genetic heritability in our sample, with an average 95% biome-explainability CI width of 32.8% (averaged over different biome-explainability levels), compared with 98.7% for a genetic heritability CI in our cohort (**Supplementary Table 11**).

We also verified that the reported results are not specific to our cohort, by using individuals from the Wellcome Trust Case Control Consortium 2 control cohorts^49^ (Methods). This sample yielded genetic heritability CIs similar to those of our cohort when subsampling to 540 individuals to match our cohort size (average 95% CI width=98.7%). Genetic heritability CIs comparable to our biome-explainability CIs were only reached when increasing the sample size to 3000 genotypes (average 95% CI width=32.0%; **Fig. 4a**; Supplementary **Table 11**).

**Figure 4:**
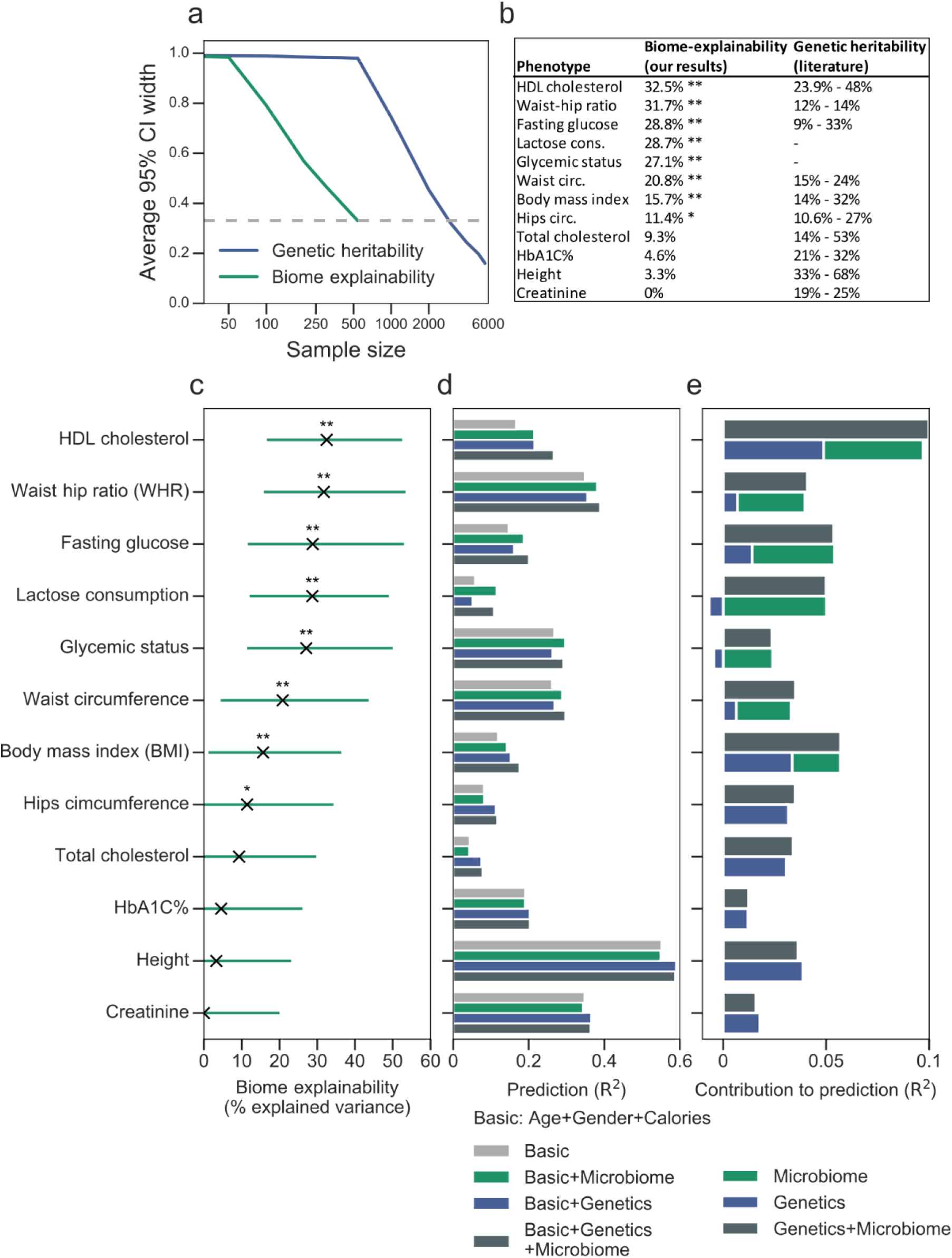
The microbiome explains a significant fraction of the variance of several human phenotypes. **(a)** Biome-explainability, defined as the phenotypic variance explained by the microbiome, can be estimated more accurately than genetic heritability. For each sample size, the values shown (y-axis) are the average 95% CI width of biome-explainability (using a kinship matrix based on bacterial genes from the present study; green curve), and of genetic heritability (using a kinship matrix of SNPs from WTCCC2 control cohorts; blue curve), computed via a parametric bootstrap and averaged over different values in the range [0,1] (Methods). Smaller CI widths indicate a greater confidence in the estimation. Biome-explainability estimation using 540 individuals provides nearly the same accuracy as genetic heritability estimation using 3,000 individuals (dashed line). Genetic heritability CIs based on our cohort of 696 individuals have a width greater than 98% for all evaluated subsets, and are omitted for clarity. **(b)** Biome-explainability estimates from our study (left column) are comparable to genetic heritability estimates from the literature (right column)^51–58^ for several human phenotypes of interest. Genetic heritability estimates are given as a range of values corresponding to the different literature estimates. **(c)** Biome-explainability estimates of several human phenotypes. Horizontal bars represent 95% confidence intervals. Glycemic status is an indicator of hyperglycemia, based on HbA1c%, fasting glucose, response to standardized meals, and data collected from continuous glucose monitors (see Methods). * FDR<0.05; ** FDR<0.01. **(d)** Phenotype prediction accuracy when using different sets of predictive features: (1) **Basic**: Age, gender, and self-reported daily average caloric, fat, protein and carbohydrates consumption; (2); **Microbiome**: Presence/absence patterns of 809,665 microbiome genes; and (3) **Genetics** - genetic factors, encoded via a polygenic risk score, which consists of obtaining effect size estimates for 554,279 SNPs from summary statistics, and then assigning a score for every individual by summing their SNPs (under a 0/1/2 encoding which corresponds to the number of minor alleles carried at each SNP), weighted by their reported effect. Prediction performance (*R*^2^) was evaluated via a 10-fold cross validation, using ridge regression with regularization of bacterial genes (Methods). **(e)** The additive contribution of microbiome and genetics to prediction performance over the basic features. The shown quantities are the difference in *R*^2^ prediction performance of a model that includes either microbiome, genetics or both, compared with a model that includes only basic features (gray bars in **Figure 4d**). Notice that the joint contribution of microbiome and genetics is similar to the sum of the individual contributions, suggesting that the contributions of microbiome and host genetics to phenotype prediction are additive and independent.

This result demonstrates that microbiome data provides a larger effective sample size than host genetics towards the task of explaining host phenotypes, indicating that there is more microbiome diversity than host genetic diversity in a given sample size. Consequently, biome-explainability estimation can be carried out with cohorts of hundreds rather than thousands of individuals.

### Microbiome explains a substantial fraction of the variance of several host phenotypes

We next estimated the biome-explainability of several host phenotypes of interest (**Table 1**). To account for genetic factors, we made use of publicly available genetic summary statistics^50^ computed from tens of thousands of individuals, by computing a polygenic risk score (PRS) for every individual and including it as a covariate in the analysis (Methods). The resulting model corresponds to the formula: phenotype = genotype effect + microbiome effect + environmental effect. An alternative approach to include genetic information in the model is to directly estimate both genetic heritability and biome-explainability in the same analysis, but as noted above, doing this for genetics requires substantially larger sample sizes than ours. We therefore restricted our analysis to phenotypes with available summary statistics and to lactose consumption, as the genetic component of lactase persistence can be largely explained via only two SNPs in European populations^43^ (Methods).

Our analysis identified several host phenotypes with a statistically significantly biome-explainability component (Method, **Fig. 4b-c**, **Supplementary Table 12**). Specifically, we found 8 of 12 traits that we tested to be significantly explained by the microbiome, with estimated biome-explainability levels of 33% for non-fasting HDL cholesterol levels, 32% for waist-hip ratio, 29% for fasting glucose, 29% for lactose consumption, 27% for glycemic status, 21% for waist circumference, and 16% for BMI.

Our estimated fraction of microbiome-explained variance for BMI and for HDL cholesterol are substantially greater than those of ref.^35^, which used a linear regression model to identify several bacterial taxa that jointly explained 4.5% of the variance of BMI and 5% of the variance of HDL cholesterol, after accounting for host genetics. We hypothesized that the difference could partly stem from our use of bacterial genes rather than bacterial taxa to explain phenotypic variance. Indeed, when using abundance of bacterial taxa rather than genes we obtained lower biome-explainability estimates (**Supplementary Fig. 4, Supplementary Table 13**), suggesting that bacterial gene abundance is more informative of human phenotypes than other microbiome characteristics.

Notably, the above biome-explainability estimates were obtained after accounting for genetic factors by including a PRS as a covariate. These biome-explainability estimates are comparable to existing SNP heritability estimates for these traits from the literature based on thousands or tens of thousands of individuals^51–58^ (**Fig. 4b**, **Supplementary Table 12**), indicating that the microbiome is an important factor associated with these traits.

### Microbiome data improves human phenotype prediction

As another comparison between host genetics and microbiome, we evaluated their ability to predict human phenotypes of interest. To this end, for each phenotype, we constructed predictive machine learning models that use bacterial gene abundances, genetic polygenic risk score (PRSs), age, sex, and daily average caloric, carbohydrates, fats and proteins consumption. The contribution of a specific data source to the phenotype can be assessed by the reduction in prediction power when excluding this data source. A small reduction indicates that either the data source is not important or that other data sources can compensate for it, whereas a large reduction indicates high predictive power. Predictions were performed with an LMM^59^ (Methods).

We found that prediction accuracy for 7 of 12 traits, including BMI, HDL cholesterol, and fasting glucose, is substantially improved when adding microbiome data on top of genetic PRSs (**Fig. 4d, Supplementary Table 14**). For example, in predicting HDL, a model without genetic or microbiome information achieves R^2^=0.17, while adding a genetic PRS improves to R^2^=0.21, and adding microbiome results in R^2^=0.27. Moreover, the contribution of both data sources is largely additive, consistent with our finding that microbiome and host genetics are mostly independent (**Fig. 4e**). We also attempted to construct predictive models by directly using our genotypes without a PRS, and obtained substantially inferior results (**Supplementary Table 15**), indicating that our sample size is not large enough to directly predict complex traits with human genomes.

Taken together, these results demonstrate that host genetics and microbiome can serve as two separate and complementary factors that explain many host phenotypes of interest. Thus, phenotype prediction can often be substantially improved by combining host genetics and microbiome data within the same prediction model.

## Discussion

In this study, we investigated the extent to which the gut microbiome composition is shaped by human genetics, using a cohort of 696 healthy Israeli individuals. Our cohort is characterized by the presence of individuals of different ancestral origins living in a relatively shared environment, and is thus particularly well suited for testing the association between microbiome and host genetics while controlling for environmental factors.

We did not detect any statistically significant microbiome-genetic associations, including associations between the microbiome and genetic ancestry, genetic kinship, or specific SNPs. From a statistical standpoint, our analysis was liberal towards trying to find a signal, as we did not apply a multiple hypothesis correction when repeating the same analysis using both metagenome and 16S data, or when using different taxonomic levels.

In contrast to the lack of association between host genetics and gut microbiome, we found significant correlations between the functional composition of gut microbiomes among individuals sharing the same household. This result corroborates previous studies showing that the human oral microbiome is dominated by household sharing^60^, and that diet reproducibly alters the gut microbiota of mice with diverse genotypes^61^. Thus, an increasing body of evidence suggests that microbiome composition is dominated by environmental factors rather than by host genetics. As several recent studies reported that the microbiome is not only stable over time^20,21^, but also resilient to some extent to perturbations like antibiotics and pathogens^62–66^, an interesting unresolved question is the extent and determinants of such stability. As a small minority of heritable microbes are unlikely to bring it about, it will be interesting to further establish which mechanisms underline microbiome stability, and which perturbations cause dysbiosis that can lead to disease susceptibility^2,67–70^.

We propose biome-explainability as a means of quantifying microbiome association with host phenotypes, show that it can be reliably estimated using metagenomic cohorts of only hundreds of individuals, and find that several phenotypes exhibit substantial biome-explainability levels in the range of 16-33%. Finally, we show that adding microbiome data to host genetics data improves prediction accuracy for several host phenotypes and that the two data sources contribute additively. We note, however, that since microbiome composition can both affect and be affected by host phenotypes, biome-explainability and microbiome-based predictions cannot be used to infer causality.

Previous studies identified heritable bacteria by observing co-occurrence among family members^22,23^, or by reporting associations between specific SNPs and bacterial taxa^23,24,26–28^. Our results are consistent with these published data, and collectively suggest that only a small number of bacteria are likely strongly heritable, and that most SNP-bacteria associations are either weak or population-dependent. These conclusions are supported by the fact that there is no overlap between the significant loci reported by any two previous studies, except for 2 SNPs in close vicinity to the LCT gene that are associated with the Bifidobacterium genus, which is likely an indirect effect as both the LCT gene and Bifidobacterium are associated with lactose consumption^43,44^. Additionally, only 5 of 211 previously reported loci replicated in our cohort. Our re-analysis of a recent twin study^23^ further estimates that the overall microbiome heritability lies between 1.9% and 8.1%. Future studies with larger sample sizes will likely identify additional heritable taxa, but are unlikely to change the overall conclusion that microbiome composition is predominantly shaped by non-genetic factors.

## Methods

### Cohort Description

This study used a cohort of individuals collected in Israel, first described in ref.^34^ Study participants were healthy individuals aged 18–70 (see full inclusion and exclusion criteria in ref.^34^). Prior to the study, participants filled medical, lifestyle, and nutritional questionnaires. All participants were monitored by a continuous glucometer (CGM) for 7 days. During that period participants were instructed to record all daily activities, including standardized and real-life meals, in real-time using their smartphones. All participants were genotyped using Illumina metabochip^71^ and provided stool samples using either a swab or an OMNIGENE-GUT (OMR-200; DNA Genotek) stool collection kit, which were metagenome-sequenced using Illumina NextSeq and HiSeq, and 16S sequenced using PCR amplification of the V3/4 region using the 515F/806R 16S rRNA gene primers followed by 500 bp paired-end sequencing Illumina MiSeq^72^. We validated that SNPs extracted from human reads in pre-filtered metagenomic sequences match SNPs extracted from the blood of their human host.

### Genotypes preprocessing and imputation

We performed stringent quality control in our initial set of 712 individuals and 712,540 SNPs. We excluded SNPs with a missingness rate >5%, Hardy-Weinberg P< 10^-9^, minor allele frequency <5%, P<0.01 for differential missingness between two batches of individuals, or a logistic regression P<10^-6^ for separation of the two batches, using PLINK^73^, yielding 554,279 SNPs for subsequent analyses. We additionally excluded individuals with >10% missing SNPs, leaving 696 individuals.

Genotypes were pre-phased using EAGLE2^74^ without a reference panel, and imputed using IMPUTE2^75^ using the 1000 genomes data set^76^ and 128 Ashkenazi Jewish individuals^77^ as reference panels. We retained only SNPs with imputation probability > 90%, and applied the filtering stages above to the imputed data, yielding 5,578,121 imputed SNPs.

### Microbial preprocessing

16S rRNA preprocessing was performed as described in our previous study^34^. For metagenome analysis, we filtered metagenomic reads containing Illumina adapters, filtered low quality reads and trimmed low quality read edges. We detected host DNA by mapping with GEM^78^ to the Human genome (hg19) with inclusive parameters, and removed human reads. We subsampled all samples to have at most 10M reads. Relative abundances from metagenomic sequencing were computed via MetaPhlAn2^79^ with default parameters. MetaPhlAn relative abundances were capped at a level of 10^−4^. We removed individuals with <10 observed species from the analysis.

When testing for association between specific taxa and specific SNPs, we log transformed the data and only used taxa present in at least 5% of individuals in our cohort, leaving us with 7/19 (remaining / total) phyla, 12/28 classes, 16/43 orders, 32/100 families, 68/229 genera and 153/673 species.

### Additional Quality Control

After all filtering stages, 665 individuals with metagenome data and 493 individuals with 16S data remained in the analysis. After additionally excluding 70 individuals with a close relative (using a kinship coefficient > 2^-11/2^, which corresponds to fifth or greater degree relatives^45^), and 21 individuals with a shared household, 573 individuals with metagenome data and 418 individuals with 16S data remained in the analysis. In the biome-explainability and phenotype prediction analyses, we used the relative abundance of genes, which required excluding individuals sequenced with only single-end reads, leaving 540 individuals.

### Gene mapping

Biome-explainability estimation and phenotype prediction were performed using bacterial gene abundances. We performed gene mapping by computing the length-normalized relative abundances of genes, obtained by similar mapping with GEM to the gene reference catalog^80^, abundance correction using an iterative algorithm based on Pathoscope^81^, and normalization to sum to 1.0. Individuals that were only sequenced with single-end reads were excluded from this analysis.

### Fasting glucose phenotyping

In the biome-explainability and phenotype prediction analyses, the fasting glucose phenotype was taken from data recorded by CGMs over a week, as described in ref.^34^ The median glucose measurement over a period of 30 minutes from self-reported wake-up time was used as a surrogate measure for fasting glucose.

### Glycemic status

For each patient we computed a quantity which we term “glycemic status” that can serve as an indicator of hyperglycemia, based on HbA1c, fasting glucose, response to standardized meals^34^, and top glucose percentiles and glucose noise as obtained from the CGM over one week. Each individual was first ranked according to each feature. The glycemic status of each individual was the median of the ranks of (1) HbA1C; (2) fasting glucose; (3) median response to standardized meals; (4) median of 90%, 95%, and 98% glucose percentiles; and (5) glucose noise. We used fasting glucose summary statistics as a surrogate measure for the PRS of this measure.

### Lactose consumption computations

We computed an estimate of average monthly lactose consumption (in grams), using a questionnaire of consumption frequency of 23 dairy products. As lactose consumption was exponentially distributed in our data, we log transformed it to induce normality for the biome-explainability and phenotype prediction analyses.

### Genetic kinship, principal components and relatedness estimation

We used PC-Relate^82^ for estimating genetic kinship and PC-AiR^37^ for genetic PC computation, as these tools are robust to the presence of relatedness and admixture. We used a filtered data set of 75,384 SNPs in approximate linkage equilibrium (r^2^<0.15), and ran an iterative estimation procedure (with the initial kinship estimates provided by KING-Robust^45^) until the PC computation converged, as described in ref.^83^ We estimated the degree of relatedness between individuals via their kinship coefficient, using the cutoffs proposed in ref.^45^ When testing kinship-ancestry associations we used the kinship matrix estimated by GCTA^84^, as the kinship matrix of PC-Relate is by definition not associated with ancestry.

### Ancestry proportions computation

For each individual, the proportion of Ashkenazi, North African, Yemen, Sephardi, Middle Eastern or unknown/other ancestry is the fraction of self-reported grandparents with that origin (**Supplementary Table 16**).

### Multiple hypothesis correction

Multiple hypothesis correction was performed via a Bonferonni correction. Note that while the Benjamini-Hochberg procedure is less conservative, it by definition cannot identify significant results when no result is significant under the Bonferonni correction. Unless stated otherwise, P values reported in the main text are corrected for multiple hypothesis testing.

### Mantel Tests

Mantel tests used were performed using Spearman correlation with 100,000 permutations. When associating a matrix to a vector, we constructed a distance matrix for the vector and then performed the test. When including covariates, they were partialed out of the matrix prior to performing the test, which is equivalent to regressing them out of its eigenvalues^39^. Unless stated otherwise, we used Euclidean distance between every pair of individuals.

### Ancestry proportions prediction

We attempted to predict ancestry proportions from microbiome composition using a variety of different techniques: Ridge regression^85^, Lasso regression^85^ and extreme gradient boosting (XGB)^86^. We used as features either the top 100 PCos of the relative abundances (using Bray-Curtis distances), the raw bacterial abundances (under various taxonomic levels) transformed to a logarithmic scale, or the PCs of the genes relative abundances, using presence/absence encoding. Prediction accuracy was measured via a 10-fold cross validation. The hyper-parameters of the methods were determined in each fold via cross validation, using only the training set of each fold.

### Canonical correlation analysis

We tested for microbiome association with host genetics via canonical correlation analysis (CCA)^40^, which takes two sets of high dimensional vectors and projects them to lower dimensional vectors that are maximally correlated. Here, the SNP vectors included all SNPs and the microbiome vector included log abundances. We regressed the covariates out of the log abundances and estimated the P value of the correlation via permutation testing. We ran CCA with either 1,2,5 or 10 components, using the implementation in Scikit-learn^87^.

### Analysis of twins data

We estimated the overall microbiome heritability, and the abundance of heritable taxa, from a data set of 3,358 twins reported by Goodrich *et al.*^23^, using the same table of operational taxonomic units (OTU) used by Goodrich *et al.* The abundance of each taxon was estimated via the sum of the relative abundances of all OTUs associated with this taxon. The overall microbiome heritability was estimated under the assumption that only taxa with heritability P value smaller than some cutoff are heritable, (using multiple cutoffs) as follows. We first assigned a heritability estimate to each OTU, given as the maximal heritability estimate among all heritable taxa associated with it. We then computed for every individual the weighted sum of the heritability estimates of OTUs associated with heritable taxa, weighted by their relative abundance. The overall microbiome heritability was estimated by averaging the computed quantity over all individuals.

### Testing for microbiome association with SNPs

We tested for associations between SNPs and microbiome β-diversity computed at various taxonomic and functional levels (phylum, class, order, family, genus, species and bacterial genes) via the MiRKAT test^42^ and with all covariates described in the main text, using the efficient implementation of RL-SKAT^88^. We note that it is also possible to use a Mantel test here, but this is computationally challenging for millions of SNPs.

### Testing for SNP-microbiome associations via the Vegan package

We repeated the technique proposed in ref. ^27^ for testing SNP associations with the microbiome via the envfit function in the vegan package^89^. We used both a regular analysis as done in ref. ^27^ and a partial analysis, which includes covariates in the model to prevent confounding. However, one disadvantage of this technique compared to our LMM-based test is that even if covariates are included in the model, the permuted SNP is assumed to be independent of these covariates, leading to an incorrect null distribution. We estimated the fraction of variance of the β-diversity matrix explained by the top ranked SNPs via the ordiR2step function in vegan.

### Testing for SNP associations with individual taxa

Association testing between individual bacteria and individual SNPs was performed using FaST-LMM^90^. We used all 665 individuals who passed quality control, including related individuals and individuals with a shared household, and controlled for these potential sources of confounding via two variance components that encode kinship (as computed via PC-Relate^82^) and household sharing (using a binary covariance matrix *G* where *G_ij_* if individuals *i* and *j* share a household and zero otherwise). When testing each SNP, we used the covariates described in the main text and a genetic kinship matrix based only on SNPs from other chromosomes to avoid proximal contamination^91^.

The abundance of bacteria present in at least 95% of individuals was encoded via the log-abundance (we excluded outlier individuals more than five standard deviations away from the mean). Otherwise, we dichotomized bacteria into presence/absence patterns and encoded the phenotype as a binary vector in order to prevent zero inflation, as it leads to a bimodal distribution (it has been demonstrated that LMMs handle binary phenotypes properly if the data was not collected via case-control sampling^92^).

### Comparing results of different studies

We compared results of previous studies by counting the number of SNPs previously reported in different studies that were in the same locus and associated with taxa belonging to the same phylum (two SNPs were considered to be in the same locus if they were <100Kb apart). We attempted to use our data to replicate results in previous studies by counting the number of SNPs with p<0.05/211, where 211 is the number of previously reported loci associated with a distinct locus and taxon. We used the closest imputed SNP when the reported SNP was not in our data.

### Relatives and household sharing tests

We tested for significant microbiome sharing among related individuals or individuals sharing a household, by comparing their average Bray-Curtis dissimilarities to that of pairs with no family relation or household sharing, via a permutation test. In each permutation, we randomly divided the combined set of all pairs into two disjoint sets while preserving the original set sizes, and asked whether the mean difference in Bray-Curtis dissimilarity between individuals in the two sets is greater than the difference observed with the real data. To prevent confounding, we only considered individuals whose stool was collected with a swab (one of the two stool collection methods). We note that standard statistical tests such as a Kolmogorov-Smirnov test cannot be employed here because of dependencies between pairs associated with the same individual.

### Computing polygenic risk scores

PRSs for traits of interest were computed using summary statistics as follows. For each phenotype, the PRS of individual *i*, 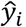, was given by 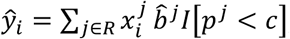, where *R* is the set of SNPs found in both the genotyping array and in the summary statistics file, 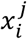 is the value of the *j^th^* SNP in the set *R* of individual *i*, 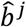 is the reported effect of SNP *j*, *p*^*j*^ is the reported univariate P value of SNP *j, I*[·] is an indicator function, and c is a P value cutoff. SNPs were normalized to have a zero mean and a unit variance (as used in the summary statistics) according to the allele frequency reported in the summary statistics. We used the original rather than the imputed set of SNPs for this task, as we empirically verified that using the imputed set of SNPs in conjunction with linkage disequilibrium pruning did not improve prediction results. The list of summary statistics used is provided in **Supplementary Table 17**.

The value of c was selected by searching over the grid [10^0^, 3 · 10^−1^,10^−1^, 3 · 10^−2^,10^−3^, …,10^−8^] and finding the value that maximizes the Spearman correlation between the true and estimated phenotypes. To prevent overfitting, we divided the data into 10 disjoint folds, estimated the value of c separately for every division of 9/10 of the folds, and then computed the PRS of the remaining individuals using the selected value. Similarly, when performing cross validation in the phenotype prediction analysis, we estimated the value of c using only individuals in the training set of each fold, and then computed the PRS for individuals in the left-out fold using this value.

### Construction of a kinship matrix based on microbiome genes

To construct a kinship matrix based on microbiome genes, we encoded the kinship of individuals i,j via 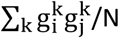, where k iterates over all genes, 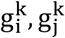 are the presence/absence indicators of gene k in individuals i and j, respectively (using a relative abundance cutoff of 10^-6^, and normalized to have a zero mean and a unit variance), and N is the number of genes (809,665). In the phenotype prediction analysis, we regressed the stool collection method out of all genes after normalizing them, to prevent confounding (this was not required in the biome explainability analysis, where this variable was included as a covariate).

### Biome-explainability estimation

Biome-explainability was estimated using GCTA^84^, using the genes-based kinship matrix. For all phenotypes (except lactose consumption), the covariates included (1) the PRS of the investigated phenotype; and (2) the covariates used in other analyses. In the analysis of lactose consumption, we replaced the PRS with the SNPs rs4988235 and rs182549, which largely explain the genetic component of lactase persistence in European populations^43^. P values were computed via RL-SKAT^88^ and confidence intervals were computed via FIESTA^93^. Outlier individuals with phenotypes more than five standard deviations away from the mean were excluded from the analysis. In several experiments reported in the Supplementary material we used a taxonomy based β-diversity matrix, which we transformed to a kinship matrix as described in ref.^42^

In the experiments analyzing the accuracy of biome-explainability estimation, the average CI width was estimated by computing the CI widths for assumed biome-explainability values in the grid [0, 0.01, 0.02, …., 1] and averaging the results.

### Analysis of data from the Wellcome Trust

We computed confidence intervals for genetic heritability estimation using 5,652 control individuals from the Wellcome trust national blood service and 1958 birth cohorts^49^. SNPs were removed with >0.5% missing data, p<0.01 for allele frequency difference between the two groups, p<0.05 for deviation from Hardy-Weinberg equilibrium, or minor allele frequency<1%. The genetic kinship matrix was computed via GCTA^84^ and confidence intervals were estimated via FIESTA^93^.

### Phenotype prediction

Phenotype prediction was performed with an LMM^59^ when including bacterial gene abundances in the model, and with linear regression otherwise. This is because LMMs reduce to linear regression in the absence of a kinship matrix. We note that LMMs are mathematically equivalent to a Ridge regression that uses the PCs of the kinship matrix as additional covariates, where only the PCs are regularized, and the regularization strength is determined via restricted maximum likelihood^94^. The LMM covariance matrix was the genes-based kinship matrix. The covariates for both models included age, sex, and daily average caloric, carbohydrates, fat and protein consumption. In some experiments we additionally included covariates for host genetic effects, represented either as a PRS (for all phenotypes except lactose consumption) or as the SNPs rs4988235, rs182549 for lactose consumption. Prediction performance was evaluated via a 10-fold cross validation. Outlier individuals with phenotypes more than five standard deviations away from the mean were excluded from all analyses.

We additionally performed experiments where we attempted to fit SNP effects directly rather than via a PRS, by including SNPs with a univariate linear regression P value <10^-5^ (estimated separately in each fold using only the training set of the fold), as additional covariates. We note that it is possible in principle to include two kinship matrices corresponding to both microbiome and genetic effects, but this was not done here due to the small sample size, which leads to inaccurate LMM training (**Fig. 4a**).

### Human Cohorts

The study was approved by Tel Aviv Sourasky Medical Center Institutional Review Board (IRB), approval numbers TLV-0658-12, TLV-0050-13 and TLV-0522-10; Kfar Shaul Hospital IRB, approval number 0-73; and Weizmann Institute of Science Bioethics and Embryonic Stem Cell Research oversight committee. Reported to http://clinicaltrials.gov/, NCT: NCT01892956.

### Data Availability

The accession number for the data reported in this paper is ENA: PRJEB11532.

## Acknowledgements

We thank the Segal and Elinav group members for fruitful discussions. S. C. thanks the Abisch-Frenkel Foundation. We thank Julia Goodrich for sharing the processed twins microbiome data with us. This study makes use of data generated by the Wellcome Trust Case Control Consortium. A full list of the investigators who contributed to the generation of the data is available from www.wtccc.org.uk. Funding for the project was provided by the Wellcome Trust under award 076113. E.S. is supported by the Crown Human Genome Center; the Else Kroener Fresenius Foundation; Donald L. Schwarz, Sherman Oaks, CA; Jack N. Halpern, New York, NY; Leesa Steinberg, Canada; and grants funded by the European Research Council and the Israel Science Foundation. D.R. received a Levi Eshkol PhD Scholarship for Personalized Medicine by the Israeli Ministry of Science.

